# PARPAL: PARalog Protein Redistribution using Abundance and Localization in Yeast Database

**DOI:** 10.1101/2025.03.04.641431

**Authors:** Brittany M. Greco, Gerardo Zapata, Rohan Dandage, Mikhail Papkov, Vanessa Pereira, François Lefebvre, Guillaume Bourque, Leopold Parts, Elena Kuzmin

## Abstract

Whole-genome duplication (WGD) events are common across various organisms however the retention and evolution of WGD paralogs is not fully understood. Quantitative measure of protein redistribution in response to the deletion of their WGD paralog provides insight into sources of gene retention. Here, we describe PARPAL (**PAR**alog **P**rotein Retribution using **A**bundance and **L**ocalization in Yeast), a web database that houses results of high-content screening and deep learning neural network analysis of the redistribution of 164 proteins reflecting how their subcellular localization and protein abundance change in response to their paralog deletion. We interrogated a total of 82 paralog pairs in two genetic backgrounds for a total of ∼3,500 micrographs of ∼460,000 cells. PARPAL also links to other studies on trigenic interactions, protein-protein interactions and protein abundance. PARPAL is available at https://parpal.c3g-app.sd4h.ca and is a valuable resource for the yeast community interested in understanding the retention and evolution of paralogs and can help researchers to investigate protein dynamics of paralogs in other organisms.

## INTRODUCTION

Gene duplication events are prevalent across the tree of life from single cell organism such as bacteria to eukaryotic multicellular complex organism such as humans (Kuzmin et al. 2022). These events result from polyploid events resulting in whole-genome duplicates (WGD) or small-scale duplicates (SSD) (Kuzmin et al. 2022). Though most duplicated genes are removed from the population, there is a significant number of paralogs that is retained through several mechanisms: neofunctionalization, whereby one paralog accumulates mutations and acquires a new function, sub-functionalization, where paralogs partition their functional domains and dosage amplification or back up compensation (Kuzmin et al. 2022).

The budding yeast, *Saccharomyces cerevisiae* underwent one round of WGD where 551 paralogs have been retained (Byrne and Wolfe 2005). Duplicated genes are thought to provide genetic robustness and thus they are an important subset of genes to study (Gu et al. 2003). Previous genetic interaction studies have shown that yeast exhibits a greater fitness defect and many trigenic interactions when both paralogs are deleted revealing a compensatory relationship between paralogs (VanderSluis et al. 2010; Kuzmin et al. 2020). On the other hand, a dependency relationship between paralogs was observed whereby a protein’s function is dependent on its paralog which often occurs in heterometric paralogs that interact physically (Diss et al. 2017).

We developed a high-content microscopy approach with single-cell resolution to understand the compensatory and dependency dynamics of paralogous proteins (Dandage et al. 2023). 328 strains harbouring a GFP-tagged protein in the wild-type or deletion background of its paralog for a total of 164 unique GFP proteins comprising 82 paralog pairs were imaged and computationally analyzed to detect a change in protein dynamics. Three different analysis methods were taken to assess protein redistribution capturing changes in subcellular localization and abundance in wild-type or deletion genetic backgrounds of their paralogs: visual inspection, protein abundance quantification and redistribution analysis. Deep neural network was used to extract 128 features and quantify redistribution. Several aspects are captured in this study: how often do proteins change their localization upon the deletion of their paralog, how often is this localization compensatory or dependent, which proteins are more likely to exhibit this localization, what drives relocalization and which features are predictive of protein redistribution. In total, 32 proteins exhibited a high redistribution score of which 30 proteins showed a change in protein abundance and 9 changed subcellular localization indicating a compensatory or dependent response (Dandage et al. 2023). Compensation was observed when a protein increased in abundance and/or changed its subcellular localization to be more similar to its paralogous protein upon the deletion of that paralog. Dependency was observed when a protein decreased in abundance and/or changed its subcellular localization to be different from its own or that of its paralogous protein which is deleted.

To enable easy access of our microscopy images of the subcellular localization and abundance changes of proteins in response to their paralog perturbation, we developed a web-accessible database called **PARPAL** (**PAR**alog **P**rotein Retribution using **A**bundance and **L**ocalization in Yeast). PARPAL currently contains ∼3,500 micrographs of ∼460,000 cells. PARPAL also links with SGD (Wong et al. 2023) and integrates Kuzmin et al. 2020 (trigenic interaction study), Diss et al. 2017 (protein-protein interaction study) and DeLuna et al. 2010 (protein abundance study) allowing for integration of multiple datasets related to yeast paralogs. This new database can be used to explore the relationship between paralog gene products in the budding yeast and differs from other databases as this catalogue of information can also be used as a resource to refer and compare findings from other studies as well as provide insight into protein dynamics of paralogs in other eukaryotic organisms.

## RESULTS AND DISCUSSION

### Microscopy data acquisition

Paralog pairs available on the PARPAL database were chosen from genes originating from WGD (Byrne and Wolfe 2005) and showed differences in their subcellular localization (Huh et al. 2003; Chong et al. 2015; Koh et al. 2015) resulting in 82 paralog pairs. In total 328 yeast query strains were constructed harbouring 164 GFP fusion proteins in two genetic backgrounds: wild-type and deletion of their paralog. Micrographs were acquired using an automated spinning disk confocal microscope with a 60x water-immersion objective (Evotec Opera, PerkinElmer). Four images per strain per replicate containing 50-100 cells were captured in a single plane totalling ∼3,500 images. These images then underwent analysis using a deep learning neural network approach for single-cell segmentation capturing ∼460,000 segmented single cells. Image analysis steps consisted of image annotation and processing, neural network architecture and training, post processing for segmentation and filtering of image data. The annotation process effectively separated individual cells from cell clumps and other imaging artifacts. Image processing allowed for the calculation of a refence background image and subtraction from all images to standardize the pixel intensity within all the images and reduce noise. A neural network was trained and used to segment cells within micrographs. A post-processing algorithm was additionally used for individual cells that did not perfectly segment previously. Finally objects smaller than 256 and larger than 8192 pixels by area were removed. Together these steps ensured that proper single cell segmentation was achieved. Images were filtered out based on a mismatch of localization between our study and a previous study (Chong et al. 2015). Images with abnormalities, artifacts, high heterogeneity as well as images with very low cell numbers were also excluded. A second deep learning neural network was used for the redistribution analysis, by extracting 128 features to capture protein abundance and subcellular localization change (Dandage 2023). Protein abundance change was also scored separately using mean GFP pixel intensities from the resulting segmented images. The workflow is shown in Figure 1.

**Figure 1.**
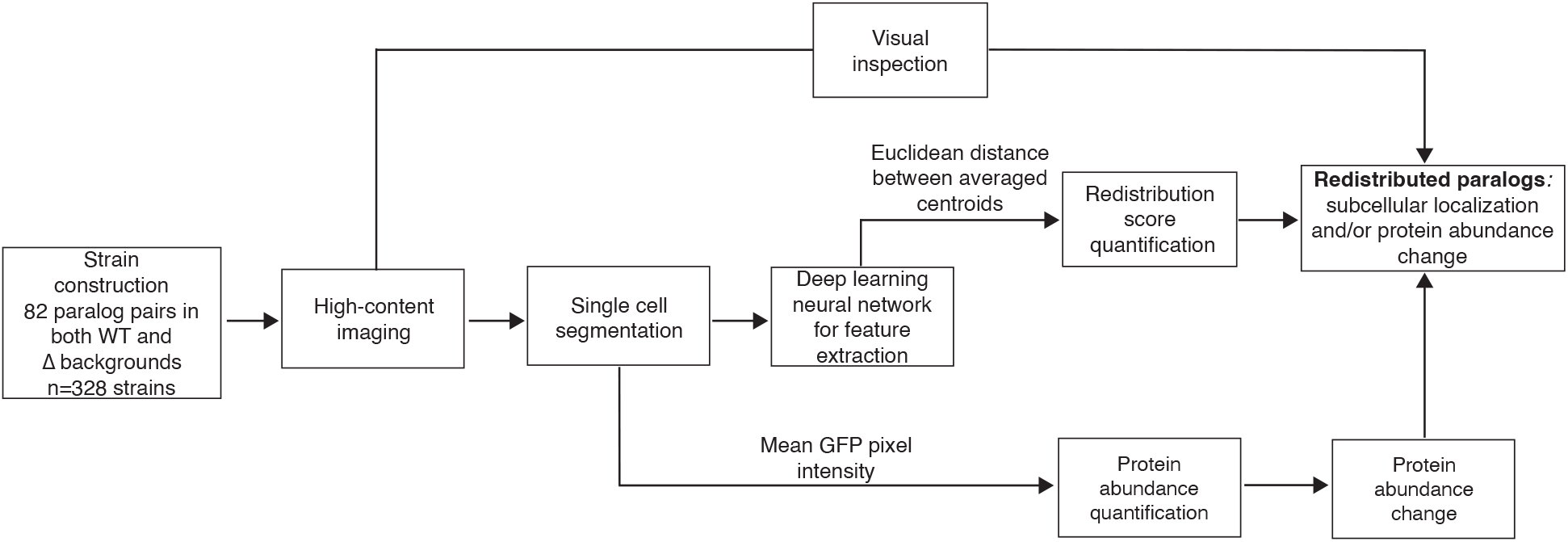
Overview of the workflow for identifying redistributed paralogous proteins using subcellular localization and protein abundance change.

### Database system construction

PARPAL was developed to facilitate the navigation of single cell paralog pair images and their corresponding protein concentration. PARPAL is a web application using shiny’s (Chang et al. 2024) framework built with python (mainly using common packages such as scikit-image, pandas, numpy and matplotlib). The app plots tables and renders images hosted by the Canadian Centre for Computational Genomics (C3G) within the S4DH’s (Secure Data for Health) secure cloud infrastructure.

### Identification of redistributed paralogs

Redistributed paralogs were identified using three different approaches: redistribution score quantification, visual inspection and independent protein abundance quantification (Figure 1). An average using an arithmetic mean across each of 128 features for all cells in three replicates was used to generate one centroid point per protein in each genetic background. The redistribution score was then defined by the Euclidean distance between the centroid for a protein in a wild-type background and the centroid of the protein in a deletion background of its paralog. The threshold for redistribution scores was calculated to be 4.73 using an AUC-ROC analysis. A Principle Component Analysis (PCA) method was used reduce dimensionality for visual representation of all 128 features. The mean GFP pixel intensity across all pixels in all the cell frames termed *abundance score* was used for protein abundance quantification in each genetic background. Abundance scores were first log-transformed and the difference between the wild-type and deletion backgrounds was calculated to generate a relative abundance change. Manual visual inspection was also used to identify cases of protein abundance and localization change. A change in protein abundance was visually seen by a decrease or increase in brightness of GFP in the microscopy image. A change in localization was denoted by a visual inspection of a change in the pattern of the GFP signal within the cells of the image. This visual inspection provided an additional level of confidence to the machine learning approach to identify the patterns associated with protein localization and abundance change in response to the paralog deletion. Together these three approaches resulted in a set of redistributed paralogs capturing subcellular localization and/or protein abundance changes (Figure 1).

### Navigating through the PARPAL database

The PARPAL web database stores redistribution scores as well as all the micrographs that were used to calculate this score. Using the pull-down menu, paralog pairs are chosen and scores will automatically appear (Figure 2A and B). Under the *Scores* tab, information such as *Paralog pair, ORF1-ORF2, Gene, ORF, redistribution score, redistribution, relative abundance change LFC, relative abundance change q-value, relative abundance change type, relocalization type* and *relocalization description* for each paralog is found (Figure 2B). *Paralog pair* is the standard name of the paralog pair, *ORF1-ORF2* is the systematic name for the paralog pair and *Gene* and *ORF* represents the systematic and standard name associated with the GFP-tagged protein, respectively. Redistribution scores higher that 4.73 indicate proteins that redistribute in response to their paralog deletion and are annotated as “true” in the *redistribution* column. Redistribution scores below the 4.73 threshold indicates that proteins which did not redistribute in response to their paralog deletion and are annotated as “false” in the *redistribution* column. Redistribution threshold was obtained based on the randomized controls and visual inspection. *Relative abundance change LFC or log-fold change* refers to the change in protein abundance of the GFP-tagged protein that was calculated from the pixel intensity values of segmented images as previously described above by comparing the abundance in the genetic background of the wild-type and deletion allele of its paralog. The *relative abundance change q-value* is the FDR-corrected p-value and was calculated using the two-sided Mann-Whitney U test and corrected for multiple testing using Benjamini/Hochberg method. *Relative abundance change type* shows whether the relative abundance change is significant if it passes the following threshold (|*log2 fold change*| *≥ 0*.*2, p < 0*.*05)* and classified into *compensation* for paralogs that increase in protein abundance or *dependency* for paralogs that decrease in protein abundance when their paralog is deleted; *ns* indicates that no significant change in protein abundance at this threshold was detected. *Relocalization description* indicates a change in subcellular localization of the GFP-protein in the absence of its paralog as evaluated by visual inspection. It reports the subcellular localization of the GFP-protein in the genetic background of wild-type allele of its paralog and the subcellular localization of GFP-protein in the genetic background of the deletion of its paralog in the following manner “subcellular localization A to subcellular localization B”. Relocalization description can also be *unclassified* if paralogs were scored to be redistributed but the subcellular localization change was not detected by visual inspection and *nan* if they were not scored as redistributed or did not change subcellular localization by visual inspection. *Relocalization type* indicates the mechanism of relocalization so that if the mechanism is said to be *compensation*, the GFP-protein changes subcellular localization to be more similar to the subcellular localization of its paralog when that paralog is deleted or *dependency*,if the GFP-protein changes subcellular localization to be different than its own subcellular localization or of its paralog in response to the deletion of that paralog. If only one protein in the paralog pair exhibits redistribution, then only values for one protein will be reported indicating that there is no redistribution for the paralog. In cases where no redistribution in either paralog pair is observed the message “No redistribution or protein abundance change” will appear under the *Scores* tab.

**Figure 2.**
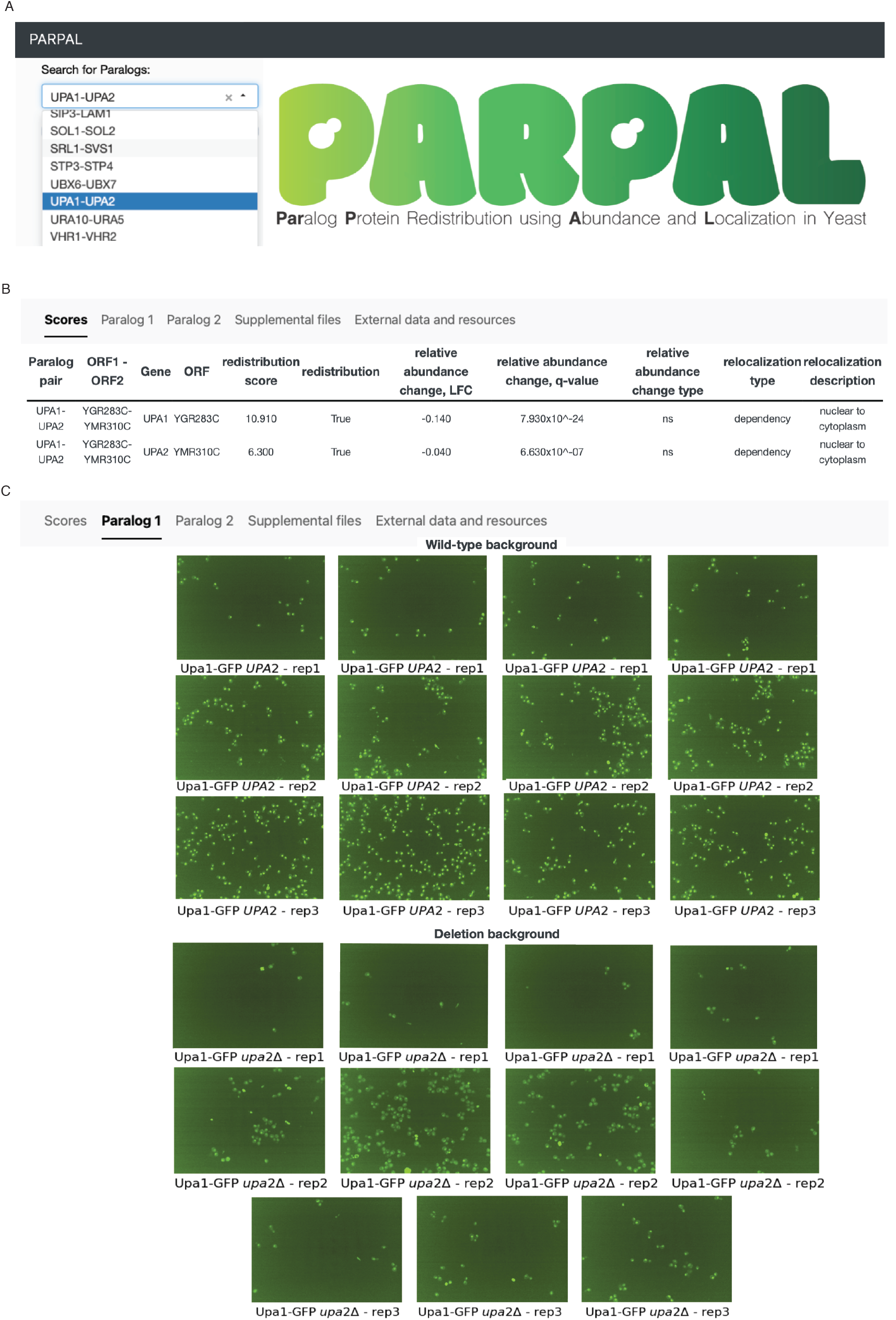
Screenshot of sample search and result page generated by PARPAL. A. A paralog pair of interest are chosen using the dropdown. B. Paralog pair, ORF1-ORF2, Gene, ORF, redistribution score, redistribution, relative abundance change, LFC, relative abundance change, q-value, relative abundance change type, relocalization type (*ns* denotes non-significant change) and relocalization description of each paralog protein (Upa1 and Upa2) are displayed in the “Scores” tab. C. In the “Paralog 1” tab, micrographs of Upa1-GFP (Paralog1-GFP) are displayed in two genetic backgrounds: wild-type *UPA2 (PARALOG2)* and deletion-background *upa2*ϕ·· (*paralog2*ϕ··). Similar to the “Paralog 1” tab, micrographs found in the “Paralog 2” tab are of Upa2-GFP (Paralog2-GFP) in both the wild-type *UPA1* (*PARALOG1)* and deletion *upa1*ϕ·· (*paralog1*ϕ··) backgrounds (not shown).

Under the *Paralog 1* and *Paralog 2* tabs, micrographs for GFP-proteins in the genetic background harboring a wild-type or deletion allele of its paralog can be found (Figure 2C). The database allows zooming in and out permitting a visual inspection of the microscopy images. Micrographs can be found in technical replicates captured from 4 fields of view as well as three biological replicates. All micrographs on PARPAL have the same microscopy settings. Some micrographs were filtered out during the quality control steps before analysis as described above. The micrographs displayed for each GFP-protein are displayed with the same threshold intensity settings to enable comparison. These parameters change across GFP-proteins and thus they should not be used to compare GFP-proteins to each other, rather to compare the same GFP protein between two different genetic backgrounds.

All supplemental information regarding all screen results housed in the PARPAL database is found in the tab called *Supplemental files*. The supplemental files consist of five tables and two datasets. Table S1 contains information regarding the yeast strains and plasmid used. Protein abundance for each paralog in each genetic background is found Table S2. Mean protein abundance as well as protein abundance for each replicate for each paralog for each condition is also included in Table S2. In Table S3 summarizes results of manual inspection. Table S4 includes all information for paralogs including redistribution, relative protein abundance changes and localization which can also be found in the *Scores* tab for paralogs that report a true statement in the redistribution column. This table additionally includes this information for protein that did not exhibit redistribution upon the deletion of its paralog. Table S5 contains furthermore information of features that may be predictors of redistribution such as protein-protein interactions, genetic interactions and colocalization of their interactors. Data S1 and Data S2 contains the scores for all 128 features from the deep neural network and protein abundance of every cell in every micrograph generated by this study, respectively.

### Exploring the protein dynamics of redistributed paralogs

The redistribution analysis revealed compensation and dependency mechanisms. Compensatory mechanisms involve the increase of protein abundance and/or the relocalization to a subcellular localization of its paralog while dependency mechanisms lead to the decrease in relative protein abundance and/or a subcellular localization different from its own and that of its paralog. An example of dependent redistribution is seen with the paralog pair, *SKN7-HMS2* (Dandage et al. 2023). Skn7 is a transcription factor needed for heat shock response to oxidative stress and osmoregulation (Raitt et al. 2000; Janiak-Spens et al. 2005) while *Hms2* is similar to a heat shock transcription factor (Engel et al. 2022). Hms2 exhibits a localization change from the nucleus to the cytoplasm upon the deletion of *SKN7*, a pattern distinct from its endogenous localization and that of its paralog (Figure 3).

**Figure 3.**
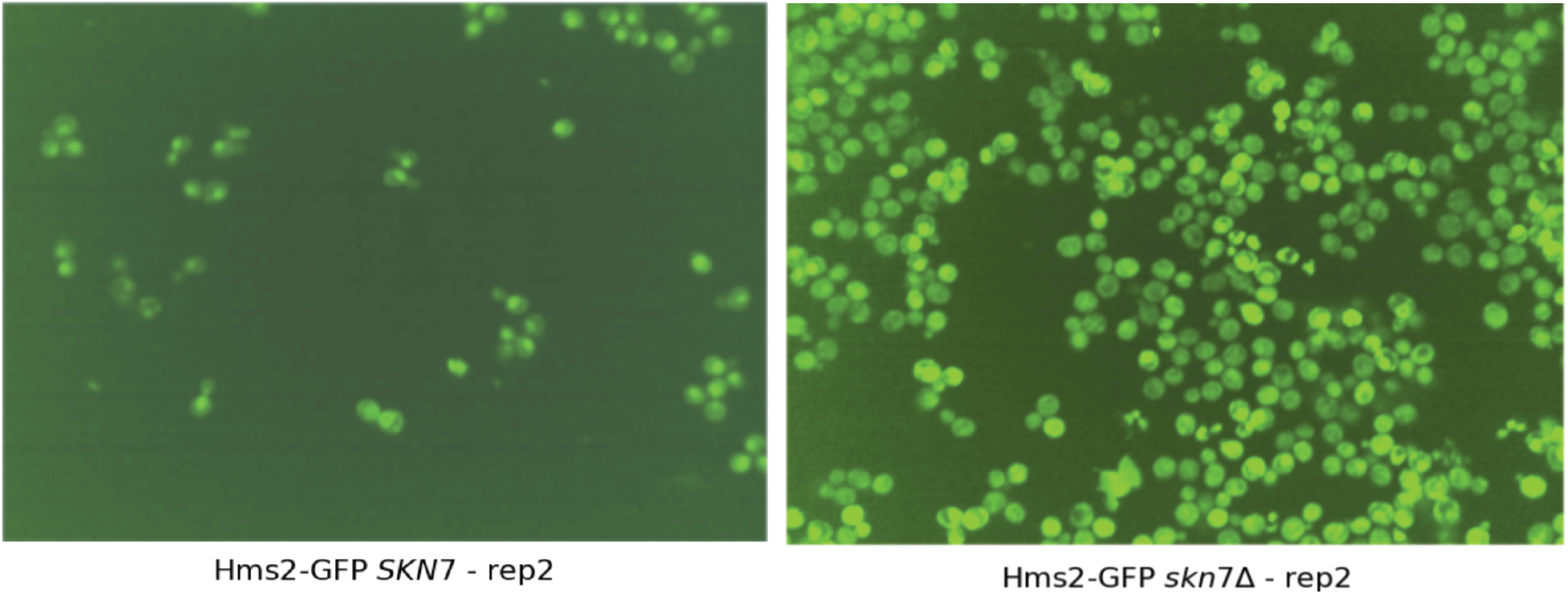
Micrographs of Hms2-GFP in the wild-type (*SKN7*) and deletion (*skn7*ϕ··) backgrounds. Micrographs of yeast cells in the wild-type background showed Hms2 localizing to the nucleus (depicted on the left) but changed localization to the cytoplasm upon the deletion of Skn7 (depicted on the right). Hms2-Skn7 is an example of a paralog pair that exhibited relocalization dependency.

### Integrating other studies into PARPAL

Genetic interactions, protein-protein interactions (PPI) and the localization of their interactors were predictive of redistribution. The PARPAL database thus includes data from our resources and studies to enhance the understanding of paralog functional divergence and redundancy, which is available in the *External data and resources* tab. Each paralog is hyperlinked to the *Saccharomyces cerevisiae* Database (SGD) (Wong et al. 2023) which provides curated and detailed information on the gene sequence and function. The database also provides a link to the digenic and trigenic interaction analysis for each paralog pair (Kuzmin et al. 2020). Low trigenic interaction fraction characterizes divergent paralogs that exhibit more paralog specific digenic interactions than trigenic interaction and was observed for *SKN7-HMS2* (Kuzmin et al. 2020). Hms2 redistributed in response to *SKN7* deletion with a dependent relocalization from the nucleus to the cytoplasm but Skn7 did not redistribute in response to *HMS2* deletion (Dandage et al. 2023) consistent with their functional divergence. High trigenic interaction fraction characterizes functionally redundant paralogs that exhibit more trigenic interactions than digenic interactions and was observed for *GGA1-GGA2* (Kuzmin et al. 2020), which encode proteins that are involved in Golgi trafficking (Zhdankina et al. 2001) . These paralogous proteins redistributed in response to each other’s deletion with Gga1 showing a compensatory relocalization from cytoplasm to Golgi in response to *GGA2* deletion and Gga2 showing a dependent protein abundance change in response to *GGA1* deletion (Dandage et al. 2023). Paralogs with a sparse number of digenic and trigenic interactions do not have a corresponding trigenic interaction fraction and are reported as ‘unclassified’ in PARPAL and include *UPA1-UPA2* (Kuzmin et al. 2020), which encode methyltransferases involved in ribosome biogenesis (Ismail et al. 2022). These paralogous proteins redistributed in response to each other’s deletion by relocalizing from the nucleus to the cytoplasm (Dandage et al. 2023).

This functional information is unavailable when examining their digenic and trigenic interactions only. Another study provided a comprehensive dataset of PPI changes of proteins in response to their paralog deletion (Diss et al. 2017). This study defines compensation when a protein expands its PPI network by gaining interactions of its paralog when that paralog is deleted and dependency when a protein loses its PPI in response to the deletion of that paralog. One of the examined paralog pairs is GSY1-GSY2 encoding glycogen synthases (Unnikrishnan et al. 2003). Despite not meeting the significance threshold for the redistribution score, which reflects the sensitivity limitation of the redistribution scoring approach, Gsy1 showed a significant protein abundance change, compensating for the loss of *GSY2* (Dandage et al. 2023). Gsy1 also showed a compensatory PPI change through an increase in its homodimer levels upon *GSY2* deletion (Diss et al. 2017). Gsy2 was classified as a redistributed protein, which compensates for the loss of *GSY1* by increasing its protein abundance (Dandage et al. 2023), although Gsy2 showed a dependent PPI with Glg1 upon *GSY1* deletion (Diss et al. 2017). Connecting the redistribution response to the changes in PPI offers mechanistic insight to the functional relationship between paralogous proteins. The third study explored responsiveness of paralogous proteins when one of the members of the pair is deleted in two types of growth conditions: rich and minimal media (DeLuna et al. 2010). Cue4 redistributed in response to *CUE1* deletion by increasing in protein abundance and changing in its subcellular localization from ER to the cytoplasm (Dandage et al. 2023). Consistent with our findings, Cue4 was upregulated upon *CUE1* deletion in rich growth media (DeLuna et al. 2010). The integration of different datasets provides insight into paralog compensation and dependency.

## CONCLUSION

We established a comprehensive dataset of protein dynamics of paralogs in the budding yeast *S. cerevisiae* which captures how proteins respond to the deletion of their paralog in terms of their subcellular localization and abundance (Dandage et al. 2023). PARPAL provides a platform to access data and visualize ∼3,500 micrographs on the redistribution of 82 paralog pairs for a total of 328 query strains and ∼460,000 cells screened. PARPAL also provides additional information of the different predictors of redistribution like genetic and protein-protein interactions by integrating other studies. Together the PARPAL database is a rich resource to provide valuable insight for other studies and expand this knowledge to high-order eukaryotes.

## ACKNOWLEDGEMENTS

We thank Leopold Parts as well as the Kuzmin lab members for helpful discussions and critical comments on the manuscript. This work was supported by Canada Research Chair program grant (CRC2021-00031 to E.K.), Canada Foundation for Innovation grant (CFI 41563 to E.K.), Natural Sciences and Engineering Research Council grant (RGPIN-2022-04674 to E.K.), Wellcome Trust (220540/Z/20/A to LP) and Revvity Inc. (VLTAT19682 to L.P.). Computing resources and data storage services were partially provided by the High-Performance Computing Center of the University of Tartu and the Digital Research Alliance of Canada.

---

**Table 1.** Summary statistics for all screens stored in PARPAL

| Genetic background | Paralog 1 | Paralog 2 | Replicates | No. proteins | No. micrographs | No. cells |
| --- | --- | --- | --- | --- | --- | --- |
| Wild-type | Paralog 1-GFP | Paralog 2 wild-type | 3 | 82 | 892 | 104,899 |
| Wild-type | Paralog 1 wild-type | Paralog 2-GFP | 3 | 82 | 853 | 96,230 |
| Deletion | Paralog 1-GFP | Paralog 2 deletion | 3 | 82 | 888 | 102,587 |
| Deletion | Paralog 1 deletion | Paralog 2-GFP | 3 | 82 | 851 | 157,235 |
|  |  |  | Total | 328 | 3,484 | 460,951 |

